# Nanodroplet-Benzalkonium Chloride Formulation Demonstrates *In Vitro* and *Ex-Vivo* Broad-Spectrum Antiviral Activity Against SARS-CoV-2 and other Enveloped Viruses

**DOI:** 10.1101/2020.11.12.377598

**Authors:** Jessie Pannu, Susan Ciotti, Shyamala Ganesan, George Arida, Chad Costley, Ali Fattom

**Author notes:** **Corresponding author**, Ali Fattom, PhD, 2311 Green Road, Suite A, Ann Arbor, MI 48105, USA.

## Abstract

The Covid-19 pandemic has highlighted the importance of aerosolized droplets inhaled into the nose in the transmission of respiratory viral disease. Inactivating pathogenic viruses at the nasal portal of entry may reduce viral loads, thereby reducing transmission and contagion. We have developed an oil-in-water nanoemulsion (*nanodroplet*) *formulation* containing the potent antiseptic 0.13% Benzalkonium Chloride (NE-BZK) which demonstrates safe and broad anti-viral activity. While The Centers for Disease Control and Prevention (CDC) have reported that BZK may have less reliable activity than ethyl alcohol against certain viruses, including coronaviruses, we have demonstrated that NE-BZK exhibits broad-spectrum, long-lasting antiviral activity with >99.99% *in vitro* killing of enveloped viruses including SARS-CoV-2, human coronavirus, RSV and influenza B. Furthermore, *in vitro* studies demonstrated that NE-BZK continues to kill >99.99% of human coronavirus even when diluted 20-fold, while 0.13% aqueous BZK solution (AQ-BZK) did not. *Ex vivo* studies of NE-BZK on human cadaver skin demonstrated persistent >99.99% killing of human coronavirus for at least 8 hours after application. AQ-BZK failed to demonstrate durable antimicrobial activity on skin over time. The repeated application of NE-BZK, twice daily for 2 weeks on to rabbit nostrils indicated safety with no irritation. These findings demonstrate that formulating BZK on the surface of proprietary nanodroplets offers a safe and effective antiviral as a significant addition to strategies to combat the spread of respiratory viral infectious diseases.

## Introduction

The nasal cavity is a primary route of entry for respiratory pathogens. Recently published studies on SARS-CoV-2 highlight the prominent role of nasal epithelium in initial infection, viral replication, and transmission.^1–3^ Other highly contagious and pathogenic viruses such as influenza, respiratory syncytial virus (RSV) and human coronavirus also enter through the nasal mucosa, causing upper respiratory infection.^4–6^ Once inoculated, viral pathogens can spread from the nose into the lower respiratory tract, causing morbidity and mortality from lower respiratory tract disease. In addition, the infected individual can spread the disease through respiratory droplets while coughing and sneezing. ^7,8^ The current Covid-19 pandemic has raised the urgent need for a broad-spectrum antiviral nasal antiseptic that can prevent infection and transmission of pathogenic respiratory viruses.

The CDC has reported varied antiviral activity of the widely used skin antiseptic AQ-BZK, including that it can be ineffective against coronavirus.^9,10^ NE-BZK is a proprietary antiseptic which incorporates the active ingredient benzalkonium chloride (0.13% BZK) in oil-in-water nanodroplets. The nanodroplets are approximately 350 nm in size and have positively charged surfaces. Given their size, they persist following application for up to 24 hours in hair follicles, sweat glands and sebaceous glands which can harbor pathogens.^11^ While these nanodroplets are small enough to penetrate these glands of the skin, they are too large to penetrate through the epidermis and are therefore not absorbed systemically. With positively charged surfaces, the nanodroplets repel each other which prevents the nanodroplets from coalescing and keeps the active ingredient BZK from inactivating through crystallization. ^12^ The surface of enveloped viruses (e.g. SARS-CoV-2, influenza, RSV) are negatively charged.^13^ Therefore, the positively charged nanodroplets in NE-BZK are attracted to these viruses, delivering the antiseptic payload directly to the surface of the pathogen where splitting and killing occurs. Ethyl alcohol products comprise over 80% of skin sanitizers available in the U.S. and have been promoted by the CDC as preferred for infection risk reduction.^9,14^ However, alcohol and water evaporate quickly after application, leading to short duration of action. Similarly, in the case of widely available aqueous 0.13% BZK antiseptics, rapid evaporation of the water component leads to breakdown of the positively charged BZK-containing micelles which determines antimicrobial activity in these formulations. In contrast, evaporation of the water component of the NE-BZK formulation leads to a residual liquid film of excipients adhering to the stratum corneum of the skin. The Poloxamer 407 co-surfactant in the nanoemulsion is a thermo-responsive hydrogel, which is bioadhesive at the surface temperature of the skin and prevents crystallization of the BZK.^15^ These advantages of NE-BZK present an attractive antiseptic formulation which can be tested for improved antiviral effect in nasal preparations. We report data from *in vitro* and *ex vivo* antiviral testing of the new formulation NE-BZK along with demonstration of *in vivo* safety of its intranasal application in rabbits.

## Results

### Broad Spectrum Antimicrobial activity of Nanoemulsion with BZK Formulation

Our experience in this field predicts that formulating cationic surfactants such as BZK in the nanoemulsion (nanodroplets) enhances its antiviral activity. Since the well-known antiseptic BZK in standard aqueous solutions has demonstrated varied activity against viruses and bacteria, including human coronavirus, we formulated BZK in oil-in-water nanodroplets. We evaluated the broad-spectrum antiviral activity of NE-BZK against various enveloped respiratory viruses. Data presented in Figure 1 shows that NE-BZK inactivated >99.99% of all tested enveloped viruses, including SARS-CoV-2. The inactivation of these viruses was achieved within 5 minutes of mixing with the nanoemulsion, indicating that the NE-BZK nanodroplets efficiently delivered the BZK to the enveloped viral particles where inactivation occurs.

**Figure 1.**
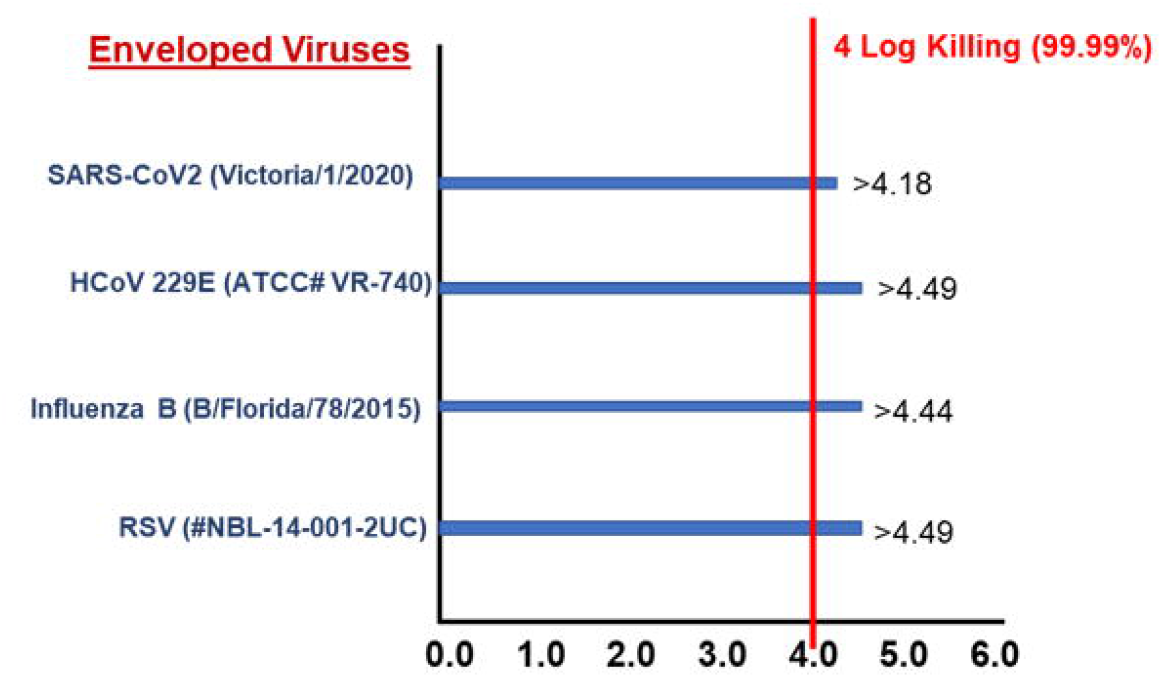
Log Reduction of Enveloped Viruses when Treated with Nanoemulsion Formulated with BZK. The antiviral effect of NE-BZK was studied by inoculating the formulation with viral particles. Following five-minute exposure, all the tested enveloped viruses demonstrated >99.99% killing, which was confirmed by plaque or TCID_50_ assay.

The results of these studies demonstrate that NE-BZK has sustained broad-spectrum antiviral activity against multiple respiratory pathogens, including SARS-CoV-2, Influenza B, and RSV.

### *In Vitro* Efficacy of Aqueous BZK in comparison to BZK formulated in Nanoemulsion against human coronavirus (HCoV229E)

To demonstrate the persistent antiviral activity of NE-BZK despite dilution, which is inherent in skin or mucosal application *in vivo,* we compared serial dilutions of NE-BZK and aqueous BZK in the *in vitro* inactivation test against human coronavirus. Three different concentrations including full-strength (0.13% BZK), 1/10 dilution (0.013% BZK) and 1/20 dilution (0.0065% BZK), were tested. Antiviral activity against human coronavirus (HCoV229E) in a time-kill study following 5 minutes exposure was evaluated. Data presented in **Table 2** demonstrate that even at highest tested dilution of 1/20, the NE-BZK formulation continued to demonstrate >99.99% killing while aqueous AQ-BZK antiviral activity was diminished with dilutions.

### Durability of NE-BZK efficacy in *ex-vivo* Human cadaver skin

Following demonstration of *in vitro* activity against human coronavirus, an ex-vivo time-kill study was performed following pre-treatment of human cadaver skin with NE-BZK (0.13% BZK) or AQ-BZK for 4 and 8 hours. As presented in **Table 3**, NE-BZK achieved >4.7 log killing of HCoV 229E at both the 4- and 8-hour time points, while AQ-BZK exhibited only 1.5 log killing at 4 hours and no detectable activity at 8 hours. These data suggest that NE-BZK could maintain its antiviral activity for a longer time due to stability of this formulation compared to AQ-BZK formulations.

### *In-vivo* Safety and Non-Irritating properties of NE-BZK product applied to Nasal Skin Surfaces

Demonstration of broad-spectrum anti-viral activity of NE-BZK makes application to the nasal cavity an option to reduce viral load, prevent colonization and lessen disease progression. Therefore, we studied the safety of multiple daily nasal administration of NE-BZK in New Zealand white rabbits. Following 2-week bi-dose intranasal application of NE-BZK, the nasal cavity was examined for skin irritation according to the method of Draize.^21^ The nasal cavity was examined for both erythema and edema and scored using the scale presented in **Table 1**, under materials and methods section. None of the animals in the treatment group (N=8) exhibited any signs of erythema or edema and were indistinguishable from the naïve group (N=4) 2 days following completing 2 weeks of twice daily treatment. Furthermore, BZK values in serum samples collected 2 hours and 2 days post application of the final dose were below the limit of detection of BZK (<0.094 μg/ml) demonstrating that this treatment localizes in the nasal cavity and does not reach the blood stream.

**Table 1.** Macroscopic Skin Irritation Grading System to Assess Erythema and Edema in Nasal Cavity.

| **Erythema Score** | **Description** |
| --- | --- |
| 0 | None, Normal color |
| 1 | Minimal, light pink, indistinct |
| 2 | Mild, bright pink/pale red, distinct |
| 3 | Moderate, bright red, distinct |
| 4 | Severe, dark red, pronounced |
| **Edema Score** | **Description** |
| 0 | None, no swelling |
| 1 | Minimal, very slight swelling, indistinct border |
| 2 | Mild, defined swelling, distinct border |
| 3 | Moderate, defined swelling, raised border (1mm) |
| 4 | Severe, pronounced swelling, raised border (>1mm) |

**Table 2.** *In vitro* log reduction of human coronavirus (HCoV 229E) treated for 5 minutes with NE-BZK verses AQ-BZK at three dilution levels

| **Product (%BZK)** | **AQ-BZK** |  | **NE-BZK** |  |
| --- | --- | --- | --- | --- |
|  | **Log Reduction** | **% Killing** | **Log Reduction** | **% Killing** |
| Full Strength (0.13%) | >4.49 | >99.99 | >4.49 | >99.99 |
| 1/10 Dilution (0.013%) | >4.49 | >99.99 | >4.49 | >99.99 |
| 1/20 Dilution (0.0065%) | <LOD* | <LOD* | >4.49 | >99.99 |

*Limit of detection (LOD) = 1*

**Table 3.** *Ex vivo* log reduction of *HCoV 229E* by Nanoemulsion Antiseptic (NE-BZK) verses an AQ-BZK at 1:10 dilutions after 4 and 8 hours post application on human skin.

| **Product**<br>**1/10 Dilution (%BZK)** | **4 Hours** | **8-Hours** |
| --- | --- | --- |
|  | **Average Log Kill\***<br>**(PFU/skin sample)** | **Average Log Kill\***<br>**(PFU/skin sample)** |
| **NE-BZK (0.013% BZK)** | >4.7 | >4.7 |
| **AQ-BZK (0.013%)** | 1.5 | <1* |

*\*Virus killing is calculated based on recovery from WFI control samples (5.70 Logs per skin sample).*
*Limit of detection (LOD) = 1 PFU*

## Discussion

NE-BZK includes in its composition a potent surfactant (0.13% BZK) that when formulated with patented oil-in-water nanodroplets, demonstrates broad-spectrum antiviral activity against enveloped viruses. The potent viral inactivation properties were demonstrated in cell culture against major respiratory viruses including two strains of coronavirus (SARS-Cov-2 and Human Coronavirus 229E), Influenza B and RSV. The nasal cavity is the primary port of entry to almost all respiratory disease-causing viruses.(reference needed) An effective nasal antiseptic that inactivates the virus at the portal of entry will be essential in preventing infection and viral transmission.^21^.

Benzalkonium chloride (BZK), a well-known effective and safe antiseptic, has demonstrated variable antiviral activity in typical aqueous formulations. In order to enhance antiviral activity and facilitate safe delivery in the nasal cavity, we formulated 0.13% BZK as the main cationic surfactant component in a new oil-in-water nanoemulsion (NE-BZK). Nanodroplets possess high adherence and residence time on epithelial tissue allowing for prolonged antiviral protection. Our data demonstrate enhanced antiviral activity vs. commonly available aqueous 0.13% BZK formulations.

The timeline of SARS-CoV-2 infection and resultant disease includes an approximately five-day incubation period before disease symptoms are displayed. High viral load and infectivity is present in infected individuals during this incubation period and 5-7 days post symptom onset. ^22, 23^ SARS-CoV-2 nasal viral load has been shown to be the most significant predictor of severe COVID-19 disease including death.^24^ Therefore, intervention aimed at reducing nasal viral load in SARS-CoV-2 infected persons would likely not only increase the probability of survival but also likely reduce viral transmission to those in contact with the infected individual. Reducing viral load and shedding in infected individuals would supplement recommended social distancing and use of face masks to mitigate the Covid-19 pandemic. A recent article by Frank et al shows the *in vitro* efficacy of povidone-iodine nasal antiseptic for rapid inactivation of SARS-CoV-2. However, this approach has major limitations including patient experience of using povidone iodine in the nose, allergic potential, and thyroid disease concerns. Povidone Iodine can become ineffective when diluting from available commercial 5% or 10% solutions and can be effective only up to 4 hours post application^25^. NE-BZK has an added advantage that safety has been previously demonstrated in humans through two different phase 1 clinical trials using the same nanoemulsion technology with a similar cationic surfactant.^26, 27^ BZK at concentrations similar to those in NE-BZK has been used in human skin products since the 1940’s and is currently marketed in numerous nasal products.(reference) Hirose et al has shown the survival of SARS-CoV-2 on skin surfaces for 9 hours^28^ which may require a more durable antiseptic activity. Our *ex-vivo* data has shown that antimicrobial effect of NE-BZK was seen even 8-12 hours post application, and even when diluting 20-fold, suggesting that application every 8 hours may be useful in protecting against disease. Our *in vivo* safety study in rabbits also demonstrated no safety issues with twice daily application of NE-BZK for two weeks, and BZK was not detected in the serum samples two-hours post application, indicating no absorption into the systemic circulation.

We believe that NE-BZK antiseptic might fulfill a role in reducing viral loads in SARS-Cov-2 infected persons, potentially reducing the risk of disease progression and increasing survival. Moreover, reduction of viral load during the isolation period may reduce transmission of the virus to households and other contacts. The attractive safety, efficacy, and easy application of NE-BZK antiviral warrants clinical evaluation as an antiviral nasal sanitizer to prevent or treat viral infections in humans at risk for viral exposure during epidemics and pandemics.

## Materials and Methods

### Nanoemulsion Formulation with BZK

The nanoemulsion formulation was prepared with 0.13% benzalkonium chloride (BZK) as the active ingredient.^16^ BZK, a quaternary ammonium compound, was chosen due to its inherent antimicrobial activity and is currently use in numerous nasal spray and skin antiseptic products. In NE-BZK, the BZK resides at the interface between the oil and water phases of the nanodroplets with the hydrophobic tail distributed in the oil core and the polar cationic head group residing at the oil-water interface.

### Virus strains used for Demonstration of Antiviral activity with NE-BZK

Broad spectrum antiviral activity of NE-BZK was tested using the following viral isolates: SARS-CoV-2 Victoria/1/2020 strain (Public Health England (PHE), Porton Down, Salisbury, UK); Human coronavirus 229E (ATCC: VR-740); Influenza B (VR-1931); and Respiratory Syncytial Virus (BlueWillow Biologics in-house strain: NBL-14-001-2UC). The virus growth media was either Minimum Essential Medium – Eagle with Earle’s BSS (MEM Eagle EBSS) from Lonza (Rochester, NY) or Dulbecco’s Modified Eagle’s Medium (DMEM) from Corning Inc (Corning, NY). The cells used in the virus studies were obtained from ATTC (Manassas, VA): Vero E6 cells (ATCC# CRL 1586) for SARS-CoV2, MRC-5 cells for HCoV229E (CCL-171 ATCC), MDCK cells for Influenza B (CCL-34 ATCC) and Vero cells for RSV (CCL-81 ATCC).

### *In Vitro* Determination of Antiviral Activity

Using the time kill procedures described in the Standard Guide ASTM E1052-11, the antiviral activity of the NE-BZK was assessed by inoculating the formulation with a suspension of viral particles (final concentration of 1.5-3.1 x10^6^ PFU/ml).^17^ At a predetermined exposure time, an aliquot was taken and neutralized to remove residual effect of NE-BZK by diluting at 1:100 dilution in cell growth media containing 1-2% FBS. A comparative inactivation of HCoV 229E by NE-BZK and AQ-BZK was also evaluated by using the test samples at full strength, 1/10 dilution, and1/20 dilution. Concentration of active virus particles was determined quantitively by plaque or TCID50 assay. Briefly, serially diluted samples were plated onto 80-90% confluent Vero E6 cells for SARS-CoV-2, MRC-5 cells for HCoV229E, MDCK for Influenza B and Vero cells for RSV. Plates were incubated for 5-7 days at 35°C for human coronaviruses and influenza B, and at 37°C for RSV, under 5% CO2. After completion of incubations, plates were fixed, stained, and counted for plaques. TCID50 was calculated by the Karber method as referenced by Lambert, based on the presence of cytopathic effect in host cells.^18^ Number of PFU recovered from the test sample was converted into log10 format and compared to an initial starting concentration to determine a log reduction.

### *Ex Vivo* Persistence of Antiviral Activity on Human Cadaver Skin following application of NE-BZK

Cryopreserved, dermatomed human cadaver abdominal skin from Caucasian donors was obtained from Science Care organ donor bank (Phoenix, AZ). The permeation and retention of each antiseptic preparation in human skin was determined using the *ex vivo* permeation technique described by Franz.^19^ Human skin was placed onto a Franz diffusion cell chamber and secured. The skin was maintained at a temperature and humidity that match typical *in vivo* conditions with a receptor phase maintained at 37°C with the water bath and magnetic stirring. The surface temperature of the skin was appropriately 32°C as determined by an Infrared surface temperature thermometer. Human skin placed onto a Franz diffusion cell chambers was dosed with either the nanoemulsion antiseptic (NE-BZK) or aqueous BZK solution (AQ-BZK) applied at a single dose of 100 μL/cm^2^. Sterile water for injection (WFI) was used as a control. At either 4 or 8 hours after the topical application, 10μL of viral particles in suspension (final concentration of 1-3 x10^5^ PFU/ml) was applied to the skin surface for a contact time of 20 minutes. The skin surface was then washed 2-3 times with 100 μL (each wash volume) of growth media. The washes were pooled and neutralized to remove residual effect of the test formulation by diluting at 1:100 dilution in cell growth media containing 1-2% FBS. Concentration of active virus particles was determined quantitively by plaque or TCID50 assay as described in the earlier section.

### *In Vivo* Rabbit Safety and Toxicology Study

The safety of NE-BZK was evaluated in New Zealand white rabbits following bi-dose nasal swab application for two weeks. The exploratory study was performed at IITRI in accordance with protocols approved by the animal care and use committee. The animals were acclimated to their designated housing for at least 7 days before the first day of dosing. The animals were randomly assigned to either naïve control or NE-BZK treatment groups with 2/sex in control group and 4/sex in NE-BZK group. The animals for NE-BZK application were held with ventral side up when intranasal swabbing was applied. The NE-BZK was applied to the inner nostril and nasal mucosa using a puritan foam tip. Care was taken that swab was applied thoroughly in ten circular motions. A clean swab was used for each nostril to prevent contamination. Two hours post last dose application and two days later, blood was collected by bleeding the central ear artery. Serum was analyzed by HPLC for BZK. The animals were sacrificed 2 days post last administration and nasal cavity was examined macroscopically and scored for erythema and edema by Draize method of scoring for dermal irritation as shown in **Table 1**.^20^

### Analysis of Serum Samples for BZK

The serum samples collected at two hours and two days post NE-BZK application was analyzed using qualified reverse phase HPLC method. The samples were extracted with three parts of acetonitrile and filtered using 0.2μm low protein binding PTFE filter before injecting into column. The samples were run on Phenomenex Luna 5μ column, using 0.04M Sodium acetate in acetonitrile as mobile phase. The samples were run at a flow rate of 2mL/ minute with 100μL injection volume. The peaks were detected using 254nm wavelength. The analytical assay detection limit was 0.094μg/mL.

### Conclusions

NE-BZK is a safe, non-irritating nasal antiseptic, which confers significant advantages over typical aqueous BZK formulations. Transmission of SARS-CoV-2 and other respiratory viruses through nasal droplets can be mitigated by inactivation of these enveloped viruses at the nasal port of entry, which may limit both the severity and the transmission of respiratory disease. NE-BZK nasal application may play a critical additional measure to prevent or reduce nasal viral load and in mitigating the Covid-19 pandemic.

## Acknowledgments

This research received no specific grant from any funding agency in the public, commercial, or not-for-profit sectors.

We are grateful to Bassam Hallis, Head of Pre-Clinical Dev & General Project Manager National Infection Service and Kevin Dyer, Senior Business Development Manager, for supporting the *in vitro* studies on SARS-CoV-2 that were conducted at the Public Health England (PHE), Porton Down *facility,* Salisbury SP4 0JG United Kingdom. We thank Sue Charlton for her excellent technical assistance in running the SARS-Cov-2 *in vitro* experiments at PHE.

We are thankful to Lakshman Caldera from BlueWilow Biologics for analyzing the serum samples from rabbit study. All of the authors whose names are listed are employees of BlueWillow Biologics.

